# Phylogenetic Identification and Functional Characterization of Orthologs and Paralogs across Human, Mouse, Fly, and Worm

**DOI:** 10.1101/005736

**Authors:** Yi-Chieh Wu, Mukul S. Bansal, Matthew D. Rasmussen, Javier Herrero, Manolis Kellis

**Affiliations:** Department of Electrical Engineering and Computer Science, Computer Science and Artificial Intelligence Laboratory, Massachusetts Institute of Technology, Cambridge, Massachusetts, USA 02139; Current address: Department of Computer Science and Engineering, University of Connecticut, Storrs, Connecticut, USA 06269; Department of Biological Statistics and Computational Biology, Cornell University, Ithaca, New York, USA 14850; European Bioinformatics Institute, European Molecular Biology Laboratory, Wellcome Trust Genome Campus, Hinxton CB10 1SD, UK; Current address: Bill Lyons Informatics Centre, UCL Cancer Institute, University College London London WC1E 6BT, UK; Broad Institute, Cambridge, Massachusetts, USA 02142

**Keywords:** phylogenomics, comparative genomics, gene duplication and loss, incomplete lineage sorting, ortholog conjecture

## Abstract

Model organisms can serve the biological and medical community by enabling the study of conserved gene families and pathways in experimentally-tractable systems. Their use, however, hinges on the ability to reliably identify evolutionary orthologs and paralogs with high accuracy, which can be a great challenge at both small and large evolutionary distances. Here, we present a phylogenomics-based approach for the identification of orthologous and paralogous genes in human, mouse, fly, and worm, which forms the foundation of the comparative analyses of the modENCODE and mouse ENCODE projects. We study a median of 16,101 genes across 2 mammalian genomes (human, mouse), 12 *Drosophila* genomes, 5 *Caenorhabditis* genomes, and an outgroup yeast genome, and demonstrate that accurate inference of evolutionary relationships and events across these species must account for frequent gene-tree topology errors due to both incomplete lineage sorting and insufficient phylogenetic signal. Furthermore, we show that integration of two separate phylogenomic pipelines yields increased accuracy, suggesting that their sources of error are independent, and finally, we leverage the resulting annotation of homologous genes to study the functional impact of gene duplication and loss in the context of rich gene expression and functional genomic datasets of the modENCODE, mouse ENCODE, and human ENCODE projects.

## 1 Introduction

Comparative analysis of complete genomes from multiple related species is a powerful technique for interpreting genes and genomes and for understanding their function and evolution (Rubin et al. 2000; Hardison 2003; Drosophila 12 Genomes Consortium 2007; Stark et al. 2007). The very premise of using model organisms to inform human biology relies on the fact that many biological processes, and the underlying genomic elements that encode them, are frequently conserved across large evolutionary distances, especially for protein-coding genes. To inform each other and human, model organism studies require a complete map of functionally-equivalent genes and processes across species, in order to facilitate mapping of biological and experimental findings across species, but such maps are extremely difficult to obtain experimentally. This need is further intensified by the human ENCODE (ENCyclopedia Of DNA Elements), mouse ENCODE, and modENCODE (model organism ENCODE) projects, to enable systematic use of the results and to further studies of functional conservation and divergence across the human, mouse, fly, and worm genomes.

The most rigorous and general method for creating a comparative genomic mapping of functionally-equivalent genes across species is through the use of phylogenetics to infer evolutionary relationships. This usually begins by first establishing a set of gene families and then studying fine-scale evolutionary events within them. Formally, a *gene family* is defined to be a group of genes such that all genes in the family are evolutionarily derived from the same ancestral gene. For any given set of species, gene families are created by first identifying all the genes in all species and then clustering them based on their sequence similarities. A pair of genes from the same gene family is said to be *homologous* to each other, and the assumption is that homologous genes are likely to have similar biological functions.

Once gene families are established, they immediately imply a coarse mapping between the genes of different species. To fine tune this mapping further, one must account for the different evolutionary events that create and shape gene families; in eukaryotic species, the most prominent evolutionary events are speciation, gene duplication, and gene loss (Ohno 1970; Demuth and Hahn 2009). Based on these events, a homologous gene pair is said to be *orthologous* if their initial divergence was the result of a speciation event, and *paralogous* if their initial divergence was the result of a gene duplication event (Fitch 1970; Koonin 2005) (Figure 1A). Distinguishing between orthologs and paralogs is important for comparative genomic studies because gene duplications are more likely to relax the evolutionary pressure on gene function (Ohno 1970; Lynch and Katju 2004; Koonin 2005; Peterson et al. 2009).

**Figure 1.**
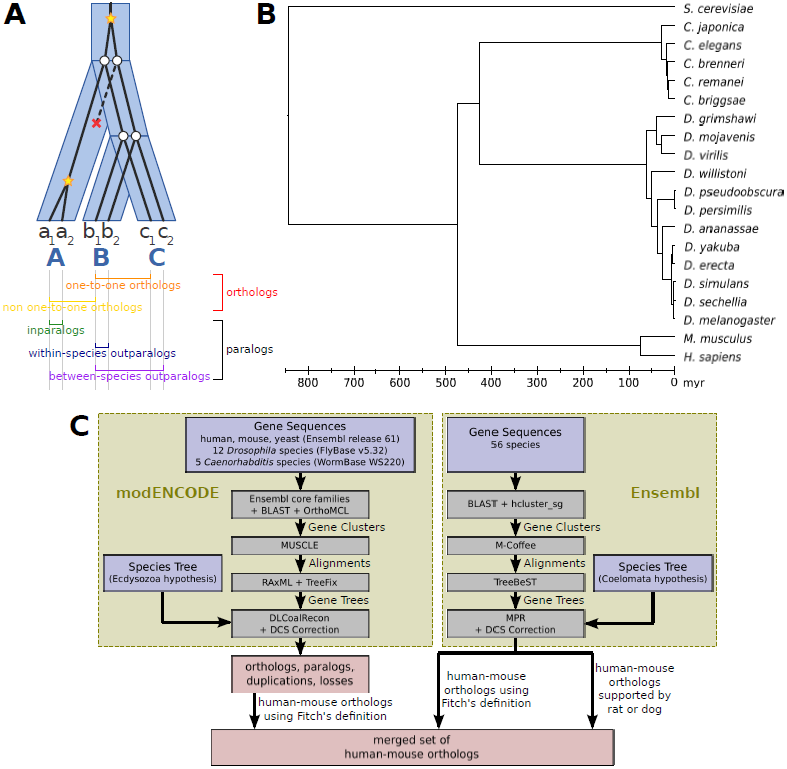
Gene family evolution. (*A*) A gene tree (black) evolves within a species tree (blue) such that incongruence between the two trees can be explained through gene duplications (yellow ‘star’) and gene losses (red ‘x’). A pair of genes is classified as *orthologs* (*paralogs*) if their most recent common ancestor is labeled as a speciation (duplication). Paralogs are further classified as *inparalogs* (*outparalogs*) if the duplication event is more (less) recent than the split between the two species. In this work, to limit the dependency on which species have been included in the analysis, paralogs are only classified as inparalogs if the duplication occurred in an extant species branch. (*B)* Phylogeny of 12 *Drosophila* species, 5 *Caenorhabditis* species, two mammals, and *S. cerevisiae* used in our analysis. (*C)* Overview of our phylogenomic pipeline. We took as input the set of all gene sequences across several species and the known species tree (blue boxes). The basic steps of the modENCODE and Ensembl pipelines are the same: (1) Gene sequences are compared across species and clustered according to sequence similarity, resulting in a set of homologous gene families. For each gene family, (2) a multiple sequence alignment is constructed, and (3) a gene tree is inferred. (4) Each gene tree is reconciled to the known species tree in order to infer orthologs, paralogs, and gene duplication and loss events, which are the pipeline outputs (red boxes). As a final step, we combined the modENCODE and Ensembl pipelines to produce a merged list of human-mouse orthologs.

Given the importance of this problem, many methods have been developed for inferring orthologs and paralogs (see, e.g., Kuzniar et al. (2008); Kristensen et al. (2011) for a review of the available methods). Approaches for ortholog/paralog inference can be generally classified into two types: those based on sequence similarities between pairs of genes in a gene family (e.g., Tatusov et al. (1997); Li et al. (2003); Alexeyenko et al. (2006); Altenhoff et al. (2011)), and tree-based methods that require a model for gene family evolution (e.g., (Dufayard et al. 2005; Li et al. 2006; van der Heijden et al. 2007; Vilella et al. 2009; Datta et al. 2009)). Methods based on sequence similarities have the advantage of being relatively fast and easy to implement, but they do not explicitly take into account the evolutionary history of the gene family and are, consequently, unable to distinguish between orthologs and paralogs based on their evolutionary definitions. On the other hand, tree-based methods explicitly reconstruct phylogenetic trees (or *gene trees*) for the gene families and reconcile them to the corresponding species phylogeny (*species tree*) to infer the evolutionary history of the gene family. For eukaryotic species, gene tree–species tree reconciliation is generally performed using a model that accounts for gene duplication and loss (Goodman et al. 1979; Page 1994), allowing for the nodes of the gene tree to be labeled as representing either a speciation or a duplication event. Given this event assignment, the task of inferring whether any given gene pair is orthologous or paralogous reduces to checking whether the most recent common ancestor of the two genes in the gene tree is a speciation event or duplication event, respectively.

While tree-based methods represent a much more principled approach to inferring orthologs and paralogs, they also suffer from several limitations in practice (Kristensen et al. 2011). One of the most crippling limitations is that gene trees can be hard to reconstruct accurately due to lack of phylogenetic signal, and inaccuracies in the gene tree translate directly into errors in inferring orthologs and paralogs (Hahn 2007). A second major handicap of existing tree-based methods is that they typically fail to account for incomplete lineage sorting (ILS) (Wakeley 1970) when reconciling gene trees with species trees, resulting in an overestimation in the number of gene duplications and losses in the gene family and incorrect inference of orthologs as paralogs (Rasmussen and Kellis 2012). Finally, for both sequence-and tree-based approaches, different methods or species samplings may yield incongruent trees and discordant orthologs and paralogs; thus, methods are needed for resolving these differences with high sensitivity and precision.

In this work, we develop a highly accurate tree-based phylogenomic pipeline for identifying orthologs and paralogs that overcomes the limitations discussed above. To address the problem of gene tree inaccuracy, we developed TreeFix (Wu et al. 2013), which, given a sequence alignment, maximum likelihood gene tree, and known species tree topology, outputs a highly accurate, error-corrected version of the gene tree. By using TreeFix to reconstruct gene trees, our method minimizes the effect of erroneous gene trees and greatly improves the accuracy of inferring orthologs and paralogs. Our approach also handles ILS effectively: the dense sampling of the fly clade and the worm clade used in modENCODE makes it especially important to distinguish ILS from gene duplication and loss (Maddison 1997; Pollard et al. 2006; Hobolth et al. 2007). To accomplish this goal, our pipeline employs DLCoal (Rasmussen and Kellis 2012), a reconciliation method that simultaneously accounts for the role of both ILS and gene duplication and loss in the evolution of a gene family through the use of an intermediate locus tree between the gene tree and species tree. Finally, we present an approach for integrating multiple ortholog sets using evidence from other species in order to benefit from the potentially complementary information available in discordant inferences.

We applied our phylogenomic pipeline to the genomes of human, mouse, twelve flies, eight worms, and yeast as part of the modENCODE and mouse ENCODE projects; the resulting orthologs and paralogs form the basis of the comparative analysis of both projects (Boyle et al. in prep; Ho et al. in prep; Gerstein et al. in prep; The mouse ENCODE Consortium et al. in prep). In this work, we present our pipeline, characterize the extent of gene tree error across these species, and demonstrate the improved accuracy of our inferences using several metrics. Furthermore, we leverage the inferred orthologs and paralogs to study the relationship between protein evolution and function, in particular, to shed light on the well-known “ortholog conjecture”, which states that orthologs are more functionally similar than paralogs.

## 2 Methods

### 2.1 Genomic sequences and species phylogeny

Genomic sequences were obtained from twelve *Drosophila* species (*D. melanogaster*, *D. simulans*, *D. sechellia*, *D. yakuba*, *D. erecta*, *D. ananassae*, *D. pseudoobscura*, *D. persimilis*, *D. willistoni*, *D. mojavensis*, *D. virilis*, *D. grimshawi*) using FlyBase (Sep 2010 release), five *Caenorhabditis* species (*C. elegans*, *C. brenneri*, *C. briggsae*, *C. japonica*, *C. remanei*) using WormBase (WS220), and two mammals (*H. sapiens*, *M. musculus*) and one outgroup species (*S. cerevisiae*) using Ensembl (release 61). We analyzed the longest protein sequence per gene (Table 1).

**Table 1.** Genome statistics.

| species | # genes<sup>a</sup> | peptide<br>length<sup>b</sup> | # families<sup>c</sup> |
| --- | --- | --- | --- |
| mammal |  |  |  |
| *H. sapiens*<sup>b</sup> | 20,477 | 540.5 | 8,561 |
| *M. musculus*<sup>b</sup> | 22,190 | 521.5 | 8,600 |
| fly |  |  |  |
| *D. melanogaster* | 13,827 | 536.7 | 7,718 |
| *D. simulans* | 15,413 | 410.5 | 8,298 |
| *D. sechellia* | 16,467 | 434.3 | 8,691 |
| *D. yakuba* | 16,077 | 468.6 | 8,415 |
| *D. erecta* | 15,044 | 484.2 | 8,349 |
| *D. ananassae* | 15,069 | 489.3 | 7,306 |
| *D. pseudoobscura* | 16,125 | 479.4 | 8,621 |
| *D. persimilis* | 16,874 | 427.3 | 8,521 |
| *D. willistoni* | 15,512 | 486.6 | 6,965 |
| *D. mojavensis* | 14,594 | 493.1 | 7,382 |
| *D. virilis* | 14,491 | 499.2 | 7,432 |
| *D. grimshawi* | 14,982 | 495.2 | 7,263 |
| worm |  |  |  |
| *C. elegans* | 20,392 | 412.6 | 7,932 |
| *C. brenneri* | 30,700 | 396.1 | 8,569 |
| *C. briggsae* | 21,967 | 397.5 | 8,087 |
| *C. japonica* | 25,870 | 324.6 | 7,511 |
| *C. remanei* | 31,471 | 399.9 | 9,089 |
| outgroup |  |  |  |
| *S. cerevisiae* | 4,010 | 546.1 | 2,676 |

<sup>a</sup> Genes for human, mouse, and yeast were filtered to non-orphaned genes from Ensembl Compara; this covers 99% of human, 96% of mouse, and 60% of yeast genes.
<sup>b</sup> Average peptide length using the longest protein sequence per gene.
<sup>c</sup> Number of non-singleton families that include at least one gene from this species.

Regarding the species phylogeny, there has been some debate on the relative grouping of mammals, *Drosophila*, and *Caenorhabditis*, with the “Ecdysozoa hypothesis” supporting mammals as the outgroup and the “Coelomata hypothesis” supporting *Caenorhabditis* as the outgroup. We used the topology consistent with the more commonly accepted Ecdysozoa hypothesis (Philippe et al. 2005; Irimia et al. 2007). Within the *Drosophila* and *Caenorhabditis* clades, we used the topologies inferred by Tamura et al. (2004) and Kiontke et al. (2004), respectively (for the full phylogeny, see Figure 1B).

Dates for the species tree were inferred using 40 one-to-one gene families (see Section 2.2) that support the (((*C. elegans*, *D. melanogaster)*, *H. sapiens*), *S. cerevisiae*) topology; these gene families were chosen as they are likely highly conserved and have not experienced duplications and losses in their histories. The alignments across these gene families were concatenated, and 10,000 sites were sampled (with replacement) from among the sites with more than two contributing sequences. We then used RAxML (Stamatakis 2006) to estimate the branch lengths given this concatenated alignment and known topology. Finally, using known dates from literature (Supplemental Table S1), remaining divergence times for the species tree were estimated using the r8s program (Sanderson 2003).

### 2.2 Phylogenomic pipeline

For our phylogenomic pipeline (Figure 1C), a reference database of sequences and core gene families was obtained from Ensembl Compara (release 61) (Vilella et al. 2009; Flicek et al. 2014). Each of the twenty genomes was BLAST (Altschul et al. 1997) against the reference database, and hits with retained if they had *e*-value ≤ 10^−5^, alignment length ≥ one-third of both query and target sequence lengths, and bitscore ≥ 80% of the bitscore for the top hit for the query sequence.

To cluster gene families, we assigned each query sequence to a core gene family based on its BLAST hits. For those genes with a hit to the reference database, most genes (97.4%) had significant similarity with only one core family. An additional set of genes (1.8%) had hits to multiple families, and the remaining sequences (0.8%) were unclustered. As some gene families may not contain a sequence from the reference database, we also ran pairwise all-vs-all BLAST comparisons for the *Drosophila* and *Caenorhabditis* proteomes, retaining hits with *e*-value ≤ 10^−5^, percent identity ≤ 60%, and alignment length ≥ one-third of both sequences. We then used OrthoMCL (Enright et al. 2002) with *I* = 1.5 to cluster genes into families.

For each gene family, we aligned the peptide sequences using MUSCLE (Edgar 2004). From this alignment, we built an initial gene tree using RAxML (Stamatakis 2006) with 100 fast bootstraps and the PROTGAMMAJTT model, then corrected for topological uncertainty using TreeFix (Wu et al. 2013), and finally accounted for possible incomplete lineage sorting using DLCoal (Rasmussen and Kellis 2012). For DLCoal, we used known parameters from literature (Supplemental Table S1).

In accordance with Fitch’s definition (Fitch 1970), we call two genes orthologs (paralogs) if their most recent common ancestor is a speciation (duplication) node. However, to improve orthology calls, we considered duplication nodes with zero consistency score (Vilella et al. 2009) as dubious and treated them as speciations. Finally, for consistency with the rest of the modENCODE and mouse ENCODE projects, we remapped Ensembl IDs to release 65.

To improve the quality of human-mouse orthologs, we also integrated results from Ensembl Compara (release 65); we used only human-mouse orthologs from Ensembl Compara as this resource focuses on vertebrates with fly and worm as outgroup species. In brief, Ensembl Compara uses hcluster_sg (http://sourceforge.net/p/treesoft/code/HEAD/tree/trunk/hcluster/) for clustering, M-Coffee (Wallace et al. 2006) for aligning the sequences, and TreeBeST (http://treesoft.sourceforge.net/treebest.shtml) for phylogenetic reconstruction. We merged human-mouse orthologs inferred using either pipeline by considering pairs of orthologous genes present in both lists. The final set of orthologs contains any pair present in both lists or present in only one of the two lists but also supported by either rat or dog, that is, a pair of genes in which the human and mouse genes are orthologous to the same genes in dog or to the same genes in rat. Lastly, the one-to-many, many-to-one and many-to-many relationships were rebuilt from the pairwise relationships.

### 2.3 Functional annotations

Enzyme commission (EC) numbers were downloaded from the ENZYME database (Jul 30, 2013).

Regulatory networks for human, *D. melanogaster*, and *C. elegans* were provided by Feizi et al. (in prep), which constructed regulatory networks by integration of multiple lines of functional and physical evidence features provided by the ENCODE and modENCODE consortia. In brief, individual networks were constructed using co-expression profiles across multiple ENCODE cell types (‘expression network’), genome-wide scans of position weight matrices (PWM) (‘motifs network’), and ChIP peak locations for multiple transcription factors (‘ChIP network’). In each network, a node represents a gene, which are either transcription factors or non-transcription factors. In the expression network, CLR (Faith et al. 2007) and GENIE3 (Huynh-Thu et al. 2010) algorithms were used to construct two expression networks, which were then merged using the Borda count (Borda 1781) network aggregation method. In the motif network, edges lead from transcription factor nodes (for which we have PWMs) to target genes where instances of the PWM are found in the 5kb promoter segment upstream of the transcription start site (TSS). In the ChIP network, edges lead from transcription factor nodes (for which ChIP peaks were called) to target genes where ChIP peaks were found in the 5kb promoter segment upstream of the TSS. Finally, the three individual networks were merged using the Borda count method.

### 2.4 Similarity metrics

For each pair of proteins within the same gene family, the sequence divergence was calculated as the fraction of paired residues (in the MUSCLE alignment) with a non-positive BLOSUM62 (Henikoff and Henikoff 1992) score. As an alternative metric, we used the age of the most recent common ancestor (MRCA), as estimated from DLCoal locus trees.

For each pair of target proteins within a regulatory network, network similarity was measured using Ruzicka’s similarity index (Ruzicka 1958) between the transcription factor families that regulate each protein. As the average similarity of random gene pairs is not equal for all genome pairs, we normalized this metric by accounting for background similarity. In particular, for all pairs of genomes (including self-pairs), we computed the background similarity by sampling with replacement 10,000 random pairs of non-homologous genes (that is, genes belonging to different families) and computing the average similarity. Relative network similarity was defined as the ratio relative to the background; throughout this work, “network similarity” always refers to this normalized score.

## 3 Results

### 3.1 Gene family properties

Using our phylogenomic pipeline, we identified 24,564 (non-singleton) gene families, including 3,108 (12.7%) mammal-specific families, of which 1,851 (59.6%) have one gene per species, 7,832 (31.9%) fly-specific families, of which 429 (5.5%) have one gene per species, and 7,295 (29.7%) worm-specific families, of which 623 (8.5%) have one gene per species. The large percentage (74.2%) of clade-specific gene families is unsurprising considering the divergence time of 474 myr between mammals and Ecdysozoa (flies and worms) and 427 myr between flies and worms.

One advantage of a tree-based approach is the ability to infer the different events responsible for gene family evolution, for example, duplications, losses, or births (generations) of novel genes. Such analysis can offer biological insights unavailable from the coarse-level clustering of gene families. For example, gene tree support, as measured by mean bootstrap support over the internal branches, is, unsurprisingly, negatively correlated with gene family size and positively correlated with alignment length (Figure 2).

**Figure 2.**
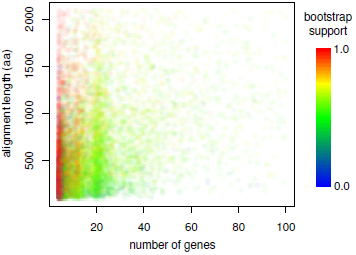
Gene tree support depends on gene family size and alignment length. For each gene family (with at least four extant genes), we determined the number of genes, alignment length, and mean bootstrap support over the internal gene tree branches.

Despite the large number of clade-specific families, we found events to be *under-represented* along the branches leading to one of the three major clades. Excluding yeast, these branches account for 53.7% of the total branch length within the species tree but contain 7,613 (31.5% of) generation events, 17,290 (24.6% of) duplication events, and 7,202 (9.3% of) loss events. We note, however, that the depletion in generation and loss events may be partially attributable to taxon sampling and annotation errors. That is, a gene family is assumed to have originated in the MRCA of all species represented in the gene family. Therefore, including other species in our phylogenomic pipeline could affect the placement of inferred events. For example, a gene must be found in both human and mouse (but no other species) for its origin to be inferred along the mammalian branch; however, if either rat or dog was included in our sampled species and a gene family was represented in either of these species, then the family could be represented in either human or mouse for its origin to be inferred along the mammalian branch. Similarly, including an animal outgroup species (as opposed to the distantly related yeast) could resolve the depletion of losses, at least partially, by moving generation events up the species tree (so that they are more ancient) followed by compensating losses along the clade branches. Finally, missing annotations could result in inferring generation events along branches within the clades rather than along branches leading to the clades.

In contrast to our finding on clade-specific events, we found events to be *over-represented* along species-specific branches. Excluding yeast, these branches account for 20.3% of the total branch length within the species tree but contain 5,658 (23.4% of), 43,186 (61.5% of), and 50,213 (64.6% of) generation, duplication, and loss events, respectively. To investigate whether this could be the result of misannotations, we focused on the well-studied reference species (human, mouse, *D. melanogaster*, *C. elegans*), which, together, account for 8.0% of the total branch length but contain 432 (1.8% of), 10,361 (14.7% of), and 9,516 (12.2% of) generation, duplication, and loss events, respectively. Note that generation events are now under-represented, and the percentages of inferred duplications and losses have reduced to 1.5–1.9× the relative branch length (compared to previously at 3.0–3.2×); this suggests that the over-representation of duplication and loss events along species-specific branches, while partially attributable to annotation errors, is likely biologically relevant.

### 3.2 Accounting for gene tree errors improves inference

A major feature of our phylogenomic pipeline is its ability to account for gene tree reconstruction error due to lack of phylogenetic signal and the confounding effects of ILS; thus, we investigated the effect of different steps in our pipeline on inference accuracy, both for the entire phylogeny and for specific clades (Table 2). Here, in addition to the number of inferred orthologs, duplications, and losses, we looked at the average duplication consistency score (DCS) (Vilella et al. 2009), which measures the support of inferred duplications.

Accounting for topological uncertainty dramatically improved inference, yielding more orthologs, fewer duplications and losses, and higher average DCS, and suggesting that many gene trees lack sufficient signal to confidently support a single topology and can be substantially improved through species-tree-informed error correction. The largest effect is seen in the fly clade, followed by the worm clade, and finally the mammalian clade; this could be indicative of the relative phylogenetic signal strength within these clades or strictly reflective of the variation in the number of species within each clade. Note also that the number of inferred losses is always more heavily reduced than the number of inferred duplications; such an observation reflects cases in which a single spurious duplication is compensated for with multiple spurious losses.

**Table 2.**
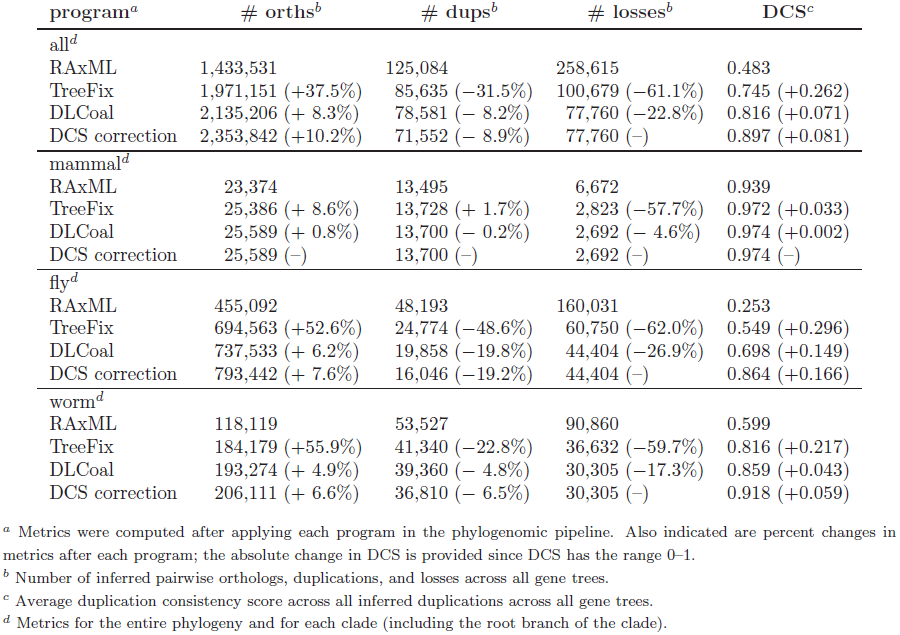
Evaluation of phylogenetic programs.

Accounting for ILS also improved inference, though to a lesser effect; the smaller effect can partially be explained since some gene tree topology errors due to ILS have already been corrected when accounting for topological uncertainty. To address ILS more directly, we also looked for incongruence between the locus tree and gene tree; that is, for each clade-specific subtree that appears in the locus tree, we asked whether the subtree appears in the gene tree. This yielded ILS rates of 0.7% for mammals, 30.1% for flies, and 19.7% for worms, which are in line with the observed differences in event inference, confirm our expectations of larger ILS rates for dense clades with shorter generation times and larger population sizes, and agree with previous reports of high ILS incidence in the fly clade (Pollard et al. 2006). Interestingly, we observe a small amount of ILS between human and mouse; it is unknown whether this is biologically relevant or an artifact of our pipeline.

### 3.3 Integrating multiple pipelines yields more accurate orthologs

A fundamental problem of phylogenomic analyses is handling differences among multiple pipelines. To address this issue, we considered inferences from Ensembl Compara to create a merged set of human-mouse orthologs, then assessed the performance of our (modENCODE) pipeline compared to these alternative ortholog sets (Table 3). Here, the modENCODE and Ensembl pipelines have their own distinct advantages: modENCODE utilizes gene tree error correction methods, which we have shown leads to improved inference, whereas Ensembl compares against many more species and may therefore have higher power to resolve orthologs versus paralogs, particularly within vertebrate species. We found that the merged set contains a slightly larger number of one-to-one orthologs while highly reducing the number of non-one-to-one orthologs; however, as the ground truth is not known for real data, we used several informative metrics to assess the quality of these inferred orthologs.

**Table 3.** Evaluation of human-mouse orthologs.

| pipeline | count<sup>a</sup> |  | e-value<sup>b</sup> |  | bootstrap<sup>b</sup> |  | GS sensitivity<sup>c</sup> |  | GS precision<sup>c</sup> |  | EC precision<sup>d</sup> |  |
| --- | --- | --- | --- | --- | --- | --- | --- | --- | --- | --- | --- | --- |
|  | 1-1 | all | 1-1 | all | 1-1 | all | 1-1 | all | 1-1 | all | 1-1 | all |
| modENCODE | 15,468 | 24,819 | 205.4 | 135.2 | 0.981 | 0.884 | 91.6% | 97.8% | 88.4% | 58.8% | 92.6% | 79.5% |
| Ensembl | 15,215 | 20,557 | 205.4 | 169.0 | 0.981 | 0.924 | 90.3% | 97.4% | 88.5% | 70.6% | 93.1% | 82.4% |
| merged | 15,736 | 19,856 | 205.2 | 176.3 | 0.980 | 0.935 | 93.5% | 97.6% | 88.6% | 73.3% | 92.9% | 82.4% |

<sup>a</sup> Comparisons against Ensembl Compara release 65.
<sup>b</sup> Median $-\log(\text{e-value})$ and mean bootstrap support.
<sup>c</sup> Percentage of 14,915 gene pairs with common gene symbol that are inferred as orthologs (sensitivity), and percentage of inferred orthologs with common gene symbol (precision).
<sup>d</sup> Percentage of inferred orthologs that share at least one enzyme commission annotation. For comparison, if we consider all gene pairs in which both genes have at least one EC annotation, then 1.2% share at least one annotation.

First, we looked at BLAST e-values between orthologs and RAxML bootstrap values for the speciation nodes relating orthologs. Note that both these metrics are based only on sequence similarity and, in particular, do not take into account other sources of information, for example, from the species tree topology. We found that the support across one-to-one orthologs is similar for all pipelines and the support across all orthologs is, unsurprisingly, lower than that for the one-to-one orthologs. However, across all orthologs, the merged set also performs better than either individual pipeline; in conjunction with the reduced counts, this suggests that merging multiple pipelines tends to remove weakly supported non-one-to-one orthologs.

Next, we examined shared annotations between inferred orthologs. As both the Human Genome Nomenclature Committee and the Mouse Genome Informatics databases manually check orthology calls when they are used to transfer gene symbols, we can regard gene pairs that share a common gene symbol as a “true” set of orthologs and use this set to determine the sensitivity and precision of our inferred orthologs. Note, however, that relying on gene symbols for orthology validation is imperfect; for example, there are instances in which symbols between non-orthologous human and mouse genes may be the same, or symbols between orthologous human and mouse genes may be different, typically for historic reasons. Furthermore, for non-one-to-one orthologs, we might expect no gene pairs to share a common gene symbol. With these short-comings in mind, we found improved one-to-one ortholog sensitivity and improved (all) ortholog precision for the merged set compared to either individual pipeline (with little change in one-to-one ortholog precision and all ortholog sensitivity); that is, the merged set shows improved overall quality. As an alternative metric, we also investigated the percentage of gene pairs that share at least one enzyme commission (EC) annotation and found improved ortholog precision for the merged set compared to the modENCODE set.

### 3.4 Regulatory networks partially support the ortholog conjecture

A major motivation of the human ENCODE, mouse ENCODE, and modENCODE projects is to provide comprehensive resources of genomic information. Therefore, in this section, we leverage this rich dataset towards understanding protein evolution and function. In particular, annotations are often transferred between homologs, and in particular, between orthologs, so we next tested this assumption that homologs share function. As a proxy for functional similarity, we used regulatory networks inferred from expression, motif, and ChIP data to compute a metric for network similarity, then normalized this metric with respect to the background. Our assumption is that if two proteins are regulated by the same transcription factors, they have similar functions. As we only have regulatory networks for human, *D. melanogaster*, and *C. elegans*, we focused on gene pairs within these species (this includes gene pairs between different species and within the same species).

We are interested in understanding both evolutionary and functional divergence of proteins and therefore compared proteins using three metrics: divergence time, sequence divergence, and network similarity (Figure 3). We observed that (1) the distribution of divergence times exhibits peaks corresponding to the speciation times between fly and worm and between mammals and Ecdysozoa, (2) the distribution of sequence divergence is centered about the middle of its range, and (3) the distribution of network similarity is centered about 1. This last finding is the most interesting as a similarity above (below) 1 implies that two genes are more (less) similar than the background; therefore, this result suggests that, as a whole, homologs are not more similar than non-homologs. However, we used as background the network similarity of non-homologs, and genes from different gene families may be functionally similar if, for instance, they are part of the same protein complex or regulatory pathway. Thus, network similarity scores below one could be a result of high background similarities, which, in turn, could be due to a high degree of interaction between gene families. Finally, we surprisingly found only moderate correlation between sequence divergence and divergence time (Pearson’s *r* = 0.322, *p* < 2.23 × 10^−308^) and weak (but significant) correlation between network similarity and divergence time (*r* = −0.083, *p* = 1.89 × 10^−251^) and between network similarity and sequence divergence (*r* = −0.098, *p* < 2.23 × 10^−308^).

**Figure 3.**
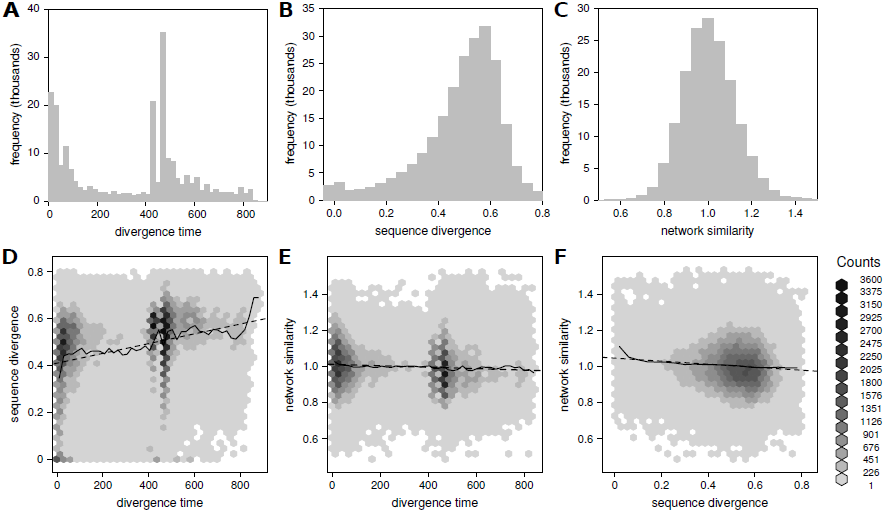
Weak to moderate correlation exists between divergence time, sequence divergence, and network similarity. (*A–C)* Histograms of the different measures, with bin sizes of 20 myr, 0.04, and 0.05, respectively. Mean and median values: 340.7, 427.0; 0.49, 0.52; 1.01, 1.00. (*D–F*) Relationships between the different measures, with the solid line depicting the mean of the dependent variable as a function of the independent variable, and the dashed line depicting the best-fit linear regression.

To further investigate the relationship between evolution and function, we considered the “ortholog conjecture”, which assumes that orthologous genes are more functionally similar than paralagous genes. However, the validity of this conjecture has recently been called into question: using GO annotations and microarray data, Nehrt et al. (2011) found that, contrary to expectation, paralogs tend to be a better predictor of function than orthologs, and they postulated that cellular context rather than sequence drives the evolution of function. However, bias in GO annotations limits their use (Thomas et al. 2012), and latter studies using GO annotations corrected for biases (Altenhoff et al. 2012) and RNA-seq data (Chen and Zhang 2012) have supported the ortholog conjecture.

To address this discrepancy, we compared the network similarity of orthologs and paralogs and found that, across the entire data set, paralogs are significantly more functionally similar than orthologs, but the difference is remarkably small (mean [median] relative network similarity for orthologs: 1.010 [0.990], for paralogs: 1.006 [1.003]; difference of 0.373% [1.236%]; Mann-Whitney test, one-tailed, *p* = 1.984 × 10^−6^). To account for variation in sequence divergence, we performed a bin-by-bin comparison of orthologs and paralogs for each level of sequence divergence and found that much of the difference in orthologs and paralogs occurs at low levels of sequence divergence (≤ 0.1); furthermore, when a significant difference is observed, paralogs tend to be more functionally similar than orthologs (Figure 4A, Supplemental Table S3).

**Figure 4.**
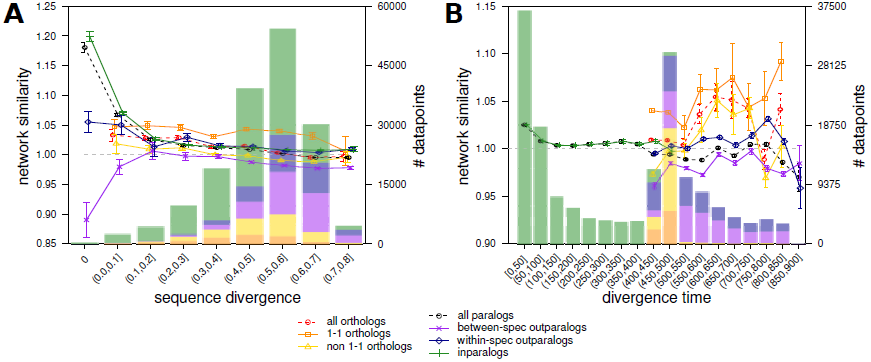
Network similarity of different types of homologs partially supports the ortholog conjecture. Mean and standard error bars are shown, and histograms represent sample density partitioned for each homology type. Orthologs with a divergence time different than the speciation time are possible due to DCS correction. Homologs are binned by (*A*) sequence divergence and (*B*) divergence time.

While the above results contradict the ortholog conjecture, orthologs and paralogs can be further sub-divided into subtypes (Figure 1A), and, by comparing the network similarity of these subtypes, we found that, ranked from the most-to-least functionally similar (as measured by network similarity) are one-to-one orthologs, inparalogs, within-species outparalogs, non-one-to-one orthologs, and between-species outparalogs (Supplemental Tables S2, S5). That is, as expected, and in agreement with the ortholog conjecture, one-to-one orthologs are the most functionally similar. As inparalogs tend to be younger than outparalogs, our finding that inparalogs are the most functionally similar subtype of paralogs is also unsurprising. Otherwise, the ranking of non-one-to-one orthologs is interesting as one might expect to find it between one-to-one orthologs and inparalogs. We believe that this result could be an artifact of species-specific annotation bias in the regulatory networks. While we hoped to minimize such bias by using a normalized network similarity metric, it is possible that some residual bias remains. The ranking of non-one-to-one orthologs could also have a biological basis – that is, with one-to-many and many-to-many orthologs, we necessarily infer one or more duplications after the speciation event that separates the orthologs, and it is possible that only one duplicate copy, if any, retains the original pre-duplicate function. Thus, for the non-one-to-one orthologs, we might expect only one pair, if any, to have similar function. It is also interesting that the network similarity of between-species outparalogs is often below the background (that is, similarity *<* 1), indicating that these paralogs are less functionally similar than non-homologs. As before, it is unknown whether this is biologically relevant or a result of high background similarities.

Surprisingly, we found little to no correlation between network similarity and sequence divergence for any homolog type (all tests that are significant with *p* < 0.001 have Spearman correlation coefficients in the range −0.087 *< ρ <* −0.028). While this contradicts the belief that sequence similarity and network similarity are expected to be highly correlated (Joshi and Xu 2007; Sangar et al. 2007), it agrees with previous studies on the ortholog conjecture (Altenhoff et al. 2012) which have shown that such is not the case.

In addition to sequence divergence, we compared the network similarity of the different homolog types at similar divergence times and found clear support for the ortholog conjecture; that is, at similar divergence times and for a wide range of times (400–850 myr), orthologs are more functionally similar than paralogs (Figure 4B, Supplemental Table S4). This stands in stark contrast to our previous finding based on sequence divergence. It also highlights the importance of accounting for dependencies when making comparisons. That is, our previous analysis conflated divergence time and rate of sequence evolution; in particular, a pair of genes that diverged recently but are found in a fast-evolving gene family (in terms of the rate of sequence evolution) could be compared against a pair of genes that diverged in ancient times but are found in a slow-evolving gene family. By utilizing divergence times, we explicitly separate these two factors and remove much of the influence of the inparalogs, the most functionally similar subtype of paralog, thereby making clear that orthologs are more similar than paralogs given the same divergence time. Interestingly, there still exists little to no correlation between relative network similarity and divergence time for any homolog type (all tests significant with *p* < 0.001 have Spearman correlation coefficients in the range −0.084 *<* ρ < 0.062).

Thus far, we have determined network similarity using modENCODE regulatory networks constructed by integration of gene expression, motif, and ChIP data. However, each of these individual networks has its own advantages and disadvantages: for example, expression data is available for most of the genome but the resulting networks are hindered by transitive edges, whereas motif and ChIP data have better sensitivity and precision at detecting regulatory interactions but are expensive and thus available for only a subset of the genome and subject to ascertainment bias that artificially inflates within-species similarities. We therefore also separately analyzed the individual networks to determine whether the features of each network have an effect on network similarity (Supplemental Figure S2). We found that many of our observations using the integrated network hold for the expression network but not for the motif or ChIP networks. In particular, using the expression network, orthologs are less functionally similar than paralogs as a whole; however, at similar levels of divergence (as measured by sequence divergence or divergence time), one-to-one orthologs are the most functionally similar, followed by inparalogs, then within-species outparalogs, then between-species outparalogs, with non-one-to-one orthologs ranking anywhere after one-to-one orthologs. Interestingly, while the expression network shows a more marked separation between the homolog types (compared to the integrated network), it also shows that orthologs are more functionally similar than paralogs at low levels of sequence divergence (≤ 0.5) but the reverse is true at high levels of sequence divergence. This threshold is important because (i) it was previously reported that functional divergence decreases dramatically above a threshold of 50% residue identity (Sangar et al. 2007) (though we do note that these results were based on problematic GO annotations), and (ii) above a sequence divergence of 0.5, mean network similarities of both orthologs and paralogs fall below the background (< 1). Therefore, it is possible that above this threshold, proteins are so diverged that network similarities become a biologically meaningless metric. As for the other networks, motif and ChIP data suffer similar problems as GO annotations, namely incomplete, biased, genome coverage, thus making these networks by themselves inappropriate for functional analysis (Thomas et al. 2012), at least not without correcting for underlying biases (Altenhoff et al. 2012). It is interesting that functional analysis using the integrated network largely agrees with analysis using the expression network; we believe this is because the expression data dominates inference in the integrated network (for example, measures for network similarity exist for 1.56× [2.52×] the number of homologs when using the expression data compared to when using motif [ChIP] data).

Finally, it has been suggested that the ortholog conjecture makes no statements about network similarity of orthologs versus paralogs. Rather, it simply states that orthologs are more likely to share function than unrelated genes. We found this interpretation to be partially supported in our analysis: using the integrated and expression networks, one-to-one orthologs and non-one-to-one homologs (the latter only if they have low sequence divergence) are more functionally similar than “unrelated” homologs.

## 4 Discussion

In this work, we have presented a highly accurate phylogenomic pipeline for inferring gene family evolution across human, mouse, fly, and worm, and using the inferred homologs and events, in combination with experimental data from the modENCODE consortium, demonstrated how joint analysis of evolutionary and functional data provides biological insight into how genes and functions arise between species.

Currently, the most popular methods for ortholog inference are based on sequence similarities or phylogenetic trees. A natural advantage of the latter approach is that, in addition to inferring orthologs (and possibly paralogs), they reconstruct gene evolutionary history. Here, we have demonstrated the utility of a tree-based phylogenomic approach towards understanding gene family evolution at the clade and species-specific levels. Furthermore, our phylogenomic pipeline infers gene trees while accounting for error due to topological uncertainty and incomplete lineage sorting; this distinguishes it from existing databases that typically ignore these issues, and, as we have demonstrated, yields dramatic improvements in event and homolog inference. In the past, the species under analysis were often separated by large evolutionary distances such that there existed sufficient phylogenetic signal (substitutions) and species were either sufficiently distant or had small populations. However, with denser sampling of the tree of life, such assumptions no longer hold, and methods that explicitly model error are needed. Indeed, our analysis clearly shows that gene tree reconstruction within dense clades can be highly inaccurate, leading to overestimation in the number of duplications and losses, underestimation in the number of orthologs, and decreased duplication consistency. Further, we have shown how our phylogenomic pipeline mitigates these effects and translates to improved phylogenetic inference across both distant and closely-related species, and we have demonstrated that performance in ortholog inference is improved by combining the results of complementary pipelines, in this case, by integrating our ortholog list for human and mouse with that of Ensembl Compara.

In addition, we have leveraged rich datasets from human, fly, and worm, as summarized through modENCODE regulatory networks, to investigate evolutionary and functional similarity and shown that that the “ortholog conjecture” is only partially supported. In particular, one-to-one orthologs are the most functionally similar subtype of homologs, and given the same divergence time, orthologs are more similar than paralogs. Remarkably, however, the difference between homolog types, though significant, is weak. In addition, under the ortholog conjecture, we might expect the functional similarity of non-one-to-one orthologs to be intermediate between one-to-one orthologs and paralogs, but in our analysis, the only consistent observation is that non-one-to-one orthologs are less functionally similar than one-to-one orthologs. We believe that this can be explained as non-one-to-one orthologs necessarily require a duplication in their joint history; if the resulting duplicates neofunctionalize or subfunctionalize, then non-one-to-one orthologs become functionally indistinguishable from paralogs. That is, the results presented here suggest that traditional analyses that distinguish only between orthologs and paralogs are insufficient. Instead, we must further characterize evolutionary relationships by the subtype of homology, and transfer of functional annotations can only be made with confidence between one-to-one orthologs. Interestingly, our analysis also highlights a vagueness in the definition of the ortholog conjecture previously brought up by Chen and Zhang (2012): the ortholog conjecture does not state whether (i) all orthologs are more functionally similar than all paralogs, or whether this holds (ii) only for those with the same divergence time or (iii) only for those with the same sequence divergence. We found support only for the second interpretation, suggesting that analyses that conflate divergence time and evolutionary rate (by using sequence divergence) provide an incomplete picture, and care must be taken to distinguish these two factors. However, we note that previous studies have supported all interpretations (Altenhoff et al. 2012; Chen and Zhang 2012), and one possibility for this discrepancy is that these other studies compared homologs across many (nine–thirteen) genomes spanning a wide range of species divergence times (36–3300 myr and 6.4–310 myr) whereas we analyze three genomes (with divergence times of 427–474 myr). Finally, we found at most a weak correlation between sequence divergence and functional divergence and between divergence time and functional divergence; this is in agreement with Altenhoff et al. (2012), which hypothesized that protein function does not evolve according to a clock.

In conclusion, this work highlights the importance of accurate phylogenomic pipelines in evolutionary analyses, presents a valuable ortholog and paralog resource for the most important animal model organisms, and provides insight into the complex relationship between evolution and function, and going forward, we believe that the use of our pipeline and resulting inferences in functional genomic and model organism studies will enable further biological insights.

## Data access

Our phylogenomic pipeline and data generated through this work are available at http://compbio.mit.edu/modencode/orthologs.

## Acknowledgements

We thank the MIT CompBio group for helpful comments, feedback, and discussions. In particular, we thank Gerald Quon, Mariana Mendoza, and Soheil Feizi for sharing their regulation networks. We are also grateful to Christophe Dessimoz for feedback on the ortholog conjecture analysis, and to various members of the ENCODE, mouse ENCODE, and modENCODE consortia for feedback on the phylogenomic pipeline and for sharing their expression, motif, and ChIP datasets.

This work was supported by the National Human Genome Research Institute (NHGRI) as part of the modENCODE project under RC2HG005639 (MK). Additional support was provided by National Institutes of Health (NIH) R01HG004037 and National Science Foundation (NSF) CAREER award 0644282 (MK), a fellowship on the MIT/Whitehead/Broad Computational Genetics Training Program training grant through the NIH (YW), and Wellcome Trust 095908, NHGRI U01HG004695, and the European Molecular Biology Laboratory (EMBL) (JH).

## References

Alexeyenko A, Tamas I, Liu G and Sonnhammer E. L. 2006. Automatic clustering of orthologs and inparalogs shared by multiple proteomes. Bioinformatics 22:e9–e15.

Altenhoff A. M, Schneider A, Gonnet G. H and Dessimoz C. 2011. Oma 2011: orthology inference among 1000 complete genomes. Nucleic Acids Res 39:D289–D294.

Altenhoff A. M, Studer R. A, Robinson-Rechavi M and Dessimoz C. 2012. Resolving the ortholog conjecture: Orthologs tend to be weakly, but significantly, more similar in function than paralogs. PLoS Comput Biol 8:e1002514–.

Altschul S. F, Madden T. L, Schaffer A. A, Zhang J, Zhang Z, Miller W and Lipman D. J. 1997. Gapped BLAST and PSI-BLAST: a new generation of protein database search programs. Nucleic Acids Res 25:3389–3402.

Borda J. 1781. Mémoire sur les Élections au scrutin. Histoire de l’Académie Royale des Sciences.

Boyle A. P, Araya C. L, Brdlik C, Cayting P, et al. (37 co-authors). in prep. Comparative analysis of regulatory information and circuits across diverse species.

Chen X and Zhang J. 2012. The ortholog conjecture is untestable by the current gene ontology but is supported by rna sequencing data. PLoS Comput Biol 8:e1002784.

Datta R. S, Meacham C, Samad B, Neyer C and Sjlander K. 2009. Berkeley phog: Phylofacts orthology group prediction web server. Nucleic Acids Res 37:W84–W89.

Demuth J. P and Hahn M. W. 2009. The life and death of gene families. BioEssays 31:29–39.

Drosophila 12 Genomes Consortium. 2007. Evolution of genes and genomes on the drosophila phylogeny. Nature 450:203–218.

Dufayard J.-F, Duret L, Penel S, Gouy M, Rechenmann F and Perrire G. 2005. Tree pattern matching in phylogenetic trees: automatic search for orthologs or paralogs in homologous gene sequence databases. Bioinformatics 21:2596–2603.

Edgar R. C. 2004. MUSCLE: multiple sequence alignment with high accuracy and high throughput. Nucleic Acids Res 32:1792–1797.

Enright A. J, Van Dongen S and Ouzounis C. A. 2002. An efficient algorithm for large-scale detection of protein families. Nucleic Acids Res 30:1575–1584.

Faith J. J, Hayete B, Thaden J. T, Mogno I, Wierzbowski J, Cottarel G, Kasif S, Collins J. J and Gardner T. S. 2007. Large-scale mapping and validation of ¡/textit¿escherichia coli¡/textit¿ transcriptional regulation from a compendium of expression profiles. PLoS Biol 5:e8–.

Feizi S, Quon G, Mendoza M, Médard M and Kellis M. in prep. Spectral network algorithms reveal conserved human, fly and worm regulatory pathways Submitted.

Fitch W. M. 1970. Distinguishing homologous from analogous proteins. Syst Biol 19:99–113.

Flicek P, Amode M. R, Barrell D, Beal K, et al. (52 co-authors). 2014. Ensembl 2014. Nucleic Acids Res 42:D749–D755.

Gerstein M. B, Rozowsky J, Yan K.-K, Wang D, et al. (95 co-authors). in prep. Comparison of 3 metazoan transcriptomes.

Goodman M, Czelusniak J, Moore G. W, Romero-Herrera A. E and Matsuda G. 1979. Fitting the gene lineage into its species lineage. a parsimony strategy illustrated by cladograms constructed from globin sequences. Syst Biol 28:132–163.

Hahn M. 2007. Bias in phylogenetic tree reconciliation methods: implications for vertebrate genome evolution. Genome Biol 8:R141.

Hardison R. C. 2003. Comparative genomics. PLoS Biol 1:e58.

Henikoff S and Henikoff J. G. 1992. Amino acid substitution matrices from protein blocks. Proceedings of the National Academy of Sciences 89:10915–10919.

Ho J. W. K, Jung Y. L, Liu T, Alver B. H, et al. (78 co-authors). in prep. Comparative analysis of metazoan chromatin architecture.

Hobolth A, Christensen O. F, Mailund T and Schierup M. H. 2007. Genomic relationships and speciation times of human, chimpanzee, and gorilla inferred from a coalescent hidden markov model. PLoS Genet 3:e7.

Huynh-Thu V. A, Irrthum A, Wehenkel L and Geurts P. 2010. Inferring regulatory networks from expression data using tree-based methods. PLoS One 5:e12776–.

Irimia M, Maeso I, Penny D, Garcia-Fernàndez J and Roy S. W. 2007. Rare coding sequence changes are consistent with ecdysozoa, not coelomata. Mol Biol Evol 24:1604–1607.

Joshi T and Xu D. 2007. Quantitative assessment of relationship between sequence similarity and function similarity. Bmc Genomics 8:222–.

Kiontke K, Gavin N. P, Raynes Y, Roehrig C, Piano F and Fitch D. H. A. 2004. *Caenorhabditis* phylogeny predicts convergence of hermaphroditism and extensive intron loss. PNAS 101:9003–9008.

Koonin E. V. 2005. Orthologs, paralogs, and evolutionary genomics. Annu Rev Genet 39:309–338.

Kristensen D. M, Wolf Y. I, Mushegian A. R and Koonin E. V. 2011. Computational methods for gene orthology inference. Brief Bioinform 12:379–391.

Kuzniar A, van Ham R. C, Pongor S and Leunissen J. A. 2008. The quest for orthologs: finding the corresponding gene across genomes. Trends Genet 24:539–551.

Li H, Coghlan A, Ruan J, Coin L. J, et al. (15 co-authors). 2006. Treefam: a curated database of phylogenetic trees of animal gene families. Nucleic Acids Res 34:D572–D580.

Li L, Stoeckert C. J and Roos D. S. 2003. Orthomcl: Identification of ortholog groups for eukaryotic genomes. Genome Res 13:2178–2189.

Lynch M and Katju V. 2004. The altered evolutionary trajectories of gene duplicates. Trends Genet 20:544– 549.

Maddison W. P. 1997. Gene trees in species trees. Syst Biol 46:523–536.

Nehrt N. L, Clark W. T, Radivojac P and Hahn M. W. 2011. Testing the ortholog conjecture with comparative functional genomic data from mammals. PLoS Comput Biol 7:e1002073–.

Ohno S. 1970. Evolution by Gene Duplication. Springer-Verlag New York.

Page R. D. M. 1994. Maps between trees and cladistic analysis of historical associations among genes, organisms, and areas. Syst Biol 43:58–77.

Peterson M. E, Chen F, Saven J. G, Roos D. S, Babbitt P. C and Sali A. 2009. Evolutionary constraints on structural similarity in orthologs and paralogs. Protein Sci 18:1306–1315.

Philippe H, Lartillot N and Brinkmann H. 2005. Multigene analyses of bilaterian animals corroborate the monophyly of ecdysozoa, lophotrochozoa, and protostomia. Mol Biol Evol 22:1246–1253.

Pollard D. A, Iyer V. N, Moses A. M and Eisen M. B. 2006. Widespread discordance of gene trees with species tree in ¡italic¿drosophila:¡/italic¿ evidence for incomplete lineage sorting. PLoS Genet 2:e173.

Rasmussen M. D and Kellis M. 2012. Unified modeling of gene duplication, loss, and coalescence using a locus tree. Genome Res 22:755–765.

Rubin G. M, Yandell M. D, Wortman J. R, Gabor G. L, et al. (56 co-authors). 2000. Comparative genomics of the eukaryotes. Science 287:2204–2215.

Ruzicka M. 1958. Anwendung mathematisch-statistischer methoden in der geobotanik (synthetische bearbeitung von aufnahmen). Biologia, Bratisl 13:647–661.

Sanderson M. J. 2003. r8s: inferring absolute rates of molecular evolution and divergence times in the absence of a molecular clock. Bioinformatics 19:301–302.

Sangar V, Blankenberg D, Altman N and Lesk A. 2007. Quantitative sequence-function relationships in proteins based on gene ontology. Bmc Bioinformatics 8:294–.

Stamatakis A. 2006. RAxML-VI-HPC: maximum likelihood-based phylogenetic analyses with thousands of taxa and mixed models. Bioinformatics 22:2688–2690.

Stark A, Lin M. F, Kheradpour P, Pedersen J. S, et al. (46 co-authors). 2007. Discovery of functional elements in 12 drosophila genomes using evolutionary signatures. Nature 450:219–232.

Tamura K, Subramanian S and Kumar S. 2004. Temporal patterns of fruit fly (*Drosophila*) evolution revealed by mutation clocks. Mol Biol Evol 21:36–44.

Tatusov R. L, Koonin E. V and Lipman D. J. 1997. A genomic perspective on protein families. Science

The mouse ENCODE Consortium, Yue F, Cheng Y, Breschi A, et al. (134 co-authors). in prep. An integrated and comparative encyclopedia of DNA elements in the mouse genome.

Thomas P. D, Wood V, Mungall C. J, Lewis S. E, Blake J. A and on behalf of the Gene Ontology Consortium. 2012. On the use of gene ontology annotations to assess functional similarity among orthologs and paralogs: A short report. PLoS Comput Biol 8:e1002386–.

van der Heijden R, Snel B, van Noort V and Huynen M. 2007. Orthology prediction at scalable resolution by phylogenetic tree analysis. Bmc Bioinformatics 8:83.

Vilella A. J, Severin J, Ureta-Vidal A, Heng L, Durbin R and Birney E. 2009. EnsemblCompara GeneTrees: Complete, duplication-aware phylogenetic trees in vertebrates. Genome Res 19:327–335.

Wakeley J. 1970. Coalescent theory: An introduction. Roberts & Company Publishers, Greenwood Village, CO.

Wallace I. M, O’Sullivan O, Higgins D. G and Notredame C. 2006. M-coffee: combining multiple sequence alignment methods with t-coffee. Nucleic Acids Res 34:1692–1699.

Wu Y.-C, Rasmussen M. D, Bansal M. S and Kellis M. 2013. Treefix: Statistically informed gene tree error correction using species trees. Syst Biol 62:110–120.

